# Castration-resistant prostate cancer cells are addicted to the high activity of cyclin-dependent kinase 2

**DOI:** 10.64898/2026.03.17.712428

**Authors:** Joyeeta Chatterjee, Alessia Marin, Shivani Yalala, Harri M Itkonen

**Affiliations:** Department of Clinical Molecular Biology, EpiGen, Institute of Clinical Medicine, University of Oslo, 1171 Oslo, Norway; Department of Biochemistry and Developmental Biology, Faculty of Medicine, University of Helsinki, 00014 Helsinki, Finland

**Keywords:** Cyclin-dependent kinase 2, cell cycle, androgen receptor, rational therapy

## Abstract

**Background:** Cyclin-dependent kinases drive the progression through the cell cycle and thereby form classical targets for cancer therapy. In prostate cancer (PC), the first line of therapy typically targets androgen receptor (AR), but it frequently leads to development of incurable form of the disease, castration-resistant PC (CRPC). Here, we sought to understand if CRPC cells are selectively addicted to a specific cell cycle kinase.

**Methods:** We used PC and CRPC patient data to evaluate transcriptional changes and modeled the responses *in vitro* using multiple models of PC, CRPC and normal cells. Development of a CDK2 inhibitor-resistant CRPC cell line, and a compound screen were used to identify chronic and acute vulnerabilities to augment the efficacy of our candidate therapy in multiple PC, CRPC and also normal cells, to assure selectivity.

**Results:** We show that the emergence of CRPC is associated with significant upregulation of cyclins that positively regulate cyclin-dependent kinase 2 (CDK2) and downregulation of CDK4 cyclins. Accordingly, CDK2-specific inhibitors and its knock down efficiently reduce proliferation of PC and CRPC cells. CDK2 inhibitor-resistant CRPC model displayed transcriptional rewiring of cell cycle regulators, characterized by a shift towards CDK4/6-dependency and increased AR-signaling. Combinatorial drug screen discovered both antagonistic and additive combinations, and we show that AR inhibitors selectively augment the efficacy of CDK2 inhibitors against PC and CRPC cells, but the combination is not toxic to normal cells.

**Conclusion:** We discovered that CRPC cells are addicted to high CDK2 activity and show that combination of CDK2 inhibitors with the currently used anti-CRPC therapies selectively augment their efficacy.

## Introduction

Prostate cancer (PC) is the most frequently diagnosed cancer and a leading cause of cancer-related mortality among men worldwide [1]. Initially, the PC is androgen-sensitive disease and can be controlled by androgen deprivation therapy (ADT). ADT targets androgen receptor (AR) signaling, and it is, at least initially, effective in most patients; however, a significant fraction develops castration-resistant PC (CRPC) [2-4]. CRPC is characterized by sustained tumor growth despite castrate levels of circulating androgens, and it is associated with poor prognosis. At this point, the disease is currently incurable, but the so-called next-generation AR-targeted agents, chemotherapy along with more experimental treatment strategies, provide extension to life-expectancy. This is the state of the disease where new treatments are needed.

All proliferating cells are dependent on the cyclin-dependent kinases (CDKs) that regulate progression through the cell cycle, which consists of two principal events: DNA replication and cell division [5, 6]. Cell cycle is controlled by the activity of cell cycle CDKs through cyclical expression of their cyclin partners. During G1 phase, mitogenic signals increase the expression of cyclin D, which binds to CDK4 and / or CDK6. The cyclin D – CDK4/6 complex mono-phosphorylates Retinoblastoma protein 1 (RB1) protein, which weakens its interaction with the E2F transcription factors. In this situation, E2F activity rises modestly leading to upregulation of cyclin E. With the accumulation of cyclin E in the G1 phase, it binds to CDK2. The cyclin E – CDK2 complex inactivates RB1 completely through its hyper-phosphorylation. This augments the expression of the E2F-driven genes required for transition from G1 to S phase and throughout the S Phase. CDK2 additionally forms complexes with cyclin E and cyclin A to stimulate replication initiation and to support S phase progression, respectively [7]. Subsequent accumulation of cyclin A/B – CDK1 complex controls mitotic entry. Cyclin A/B – CDK1 complex activates APC/CDC20 ubiquitin ligase, which promotes mitotic exit through targeted cyclin-degradation, thereby assuring, in part, that the DNA is replicated only once within the S Phase [8]. In mammalian cells, CDK1 can drive progression through the cell cycle in the absence of CDK2, CDK4 and CDK6, which in part has motivated development of selective inhibitors against the three non-essential kinases that cancer cells can have acquired dependency of [9, 10].

In cancer, CDKs are hyper-activated due to increased expression of CDKs themselves, cyclins and / or decreased expression of the endogenous CDK inhibitory proteins [11, 12]. This results in uncontrolled transition throughout the different cell cycle-phases and may open cancer cell-selective vulnerabilities. In breast cancer, the notion that Cyclin D1 is over-expressed in the tumor cells and also required for development of breast cancer, has motivated clinical trials and sub-sequent approval of CDK4/6 inhibitors against breast cancer [13-15]. CDK4/6 inhibitors aim to arrest the cells to G1-phase of the cell-cycle and suppress tumor proliferation, while sparing the normal non-cancerous tissues, which is further accomplished by targeting cancer cell-specific anomaly, dependency on the estrogen receptor. The success in breast cancer has motivated clinical trials also in PC; however, the CDK4/6 inhibitors have failed to control the disease and their development has been largely discontinued (see, for example NCT05999968, NCT02905318, NCT05617885 and NCT04408924) [16, 17]. Given that breast cancer cells are addicted to CDK4/6 activity, it raises the possibility that PC cells are dependent on other cell cycle CDKs.

Increased expression of Cyclin E (CCNE) has been proposed as a biomarker for acquired dependency on CDK2, which can guide treatment selection for the selective CDK2-targeted therapies [18]. BLU-222 and Tagtociclib (PF-07104091) are investigational, potent and highly selective inhibitors of CDK2 [19]. In *CCNE1*-amplified endometrial cancer cells, BLU-222 disrupts the RB-signaling, leading to G1 cell cycle-arrest [20]. Further preclinical *in vitro* and *in vivo* studies suggest that combining BLU-222 with other therapies provides additional benefit in the *CCNE1*-altered endometrial cancers [20]. Currently, Phase I/II clinical trials are evaluating Tagtociclib in ovarian cancer and non-small cell lung cancer to assess safety and early efficacy signals (NCT05262400, NCT04553133).

Here, we sought to establish if altered CDK2 signaling is associated with emergence of the CRPC-phenotype and if CDK2 inhibitors can control proliferation of CRPC cells. We show that development of metastatic CRPC results in significant upregulation of the E-type cyclins and concomitant downregulation of the D-type cyclins. CDK2 inhibition suppresses proliferation of CRPC cells but fails to eliminate them. We develop CDK2 inhibitor-resistant CRPC model and map potential combinatorial toxicities with clinically used CRPC-therapies in combination with BLU-222 and Tagtociclib. Our data reveal that resistance to CDK2 inhibition results in acquired dependency on CDK4/6. In addition, resistant cells hyper-activate AR-signaling, and, accordingly, the combination of anti-androgens with CDK2 inhibitors is lethal to CRPC cells. This study suggests rational for clinical trial for assessing CDK2 inhibitors in CRPC or hormone-dependent metastatic PC.

## Methods

### Cell culture, compounds, preparation of cell lysates and isolation of mRNA

LNCaP, C4-2, 22RV1, and RWPE-1 cell lines were obtained from the American Tissue Culture Collection, while PNT1 cells were obtained from Sigma; all were maintained in the RPMI media supplemented with 10% fetal bovine serum (FBS). RWPE-1 cells were maintained in keratinocyte serum free media. For androgen-deprivation experiments, 10 % charcoal-stripped serum in phenol red-free RPMI was used. In all of the experiments, cells were allowed to adhere to the plate for at least one day before starting the treatments. BLU-222, tagtociclib, docetaxel, cabazitaxel, carboplatin, cisplatin, etoposide, olaparib, rucaparib, everolimus, CX-5461, enzalutamide and darolutamide were obtained from MedChemExpress. For radiation experiments, the cells were exposed to a single dose of radiation (4 Gy).

For cell lysate preparation we followed a previous protocol and performed every step at 4°C [21]. In brief, cells were washed once with PBS, lysed with RIPA buffer (10 mmol/L Tris-HCl, pH 8.0, 1 mmol/L EDTA, 0.5 mmol/L EGTA, 1% Triton X-100, 0.1% sodium-deoxycholate, 0.1% SDS, 140 mmol/L NaCl), and supplemented with freshly added protease and phosphatase inhibitors (MedChemExpress), incubated for 30 minutes, centrifuged at 15 000 rpm for 10 minutes, and the supernatant was collected. The protein concentration was measured using the bicinchoninic acid (BCA) assay after the supernatant was collected. Antibodies were obtained from: Cell Signaling Technologies: CDK2 (2546), Proteintech: p53 (60283-2-Ig), and Abcam: Actin (ab49900). Horseradish peroxidase-conjugated secondary antibodies against cognate species were used to identify primary antibodies.

For knockdown experiments, transfection of Silencer® Select siRNAs against CDK2 was achieved using RNAiMax (ThermoFisher Scientific: Catalog # 4427038, IDs: s204 and 206). For the validation of knockdown, cell lysate was collected after 72 hours of transfection for western blot analysis as described above.

### Proliferation assays and knockdown

Cells were treated one day after the plating, and cell viability was assessed after four days of treatment using the CellTiter-Glo® 2.0 assay (Promega) according to the manufacturer’s instructions. Colony-formation assay was performed as previously reported [22]. In brief, cells were allowed to grow and adhere for 24 hours prior to treatment. After one week of the treatment, the cells were washed with PBS buffer and fixed onto the plates using ice-cold 70% methanol for 2 minutes, followed by 100% methanol for 10 minutes. Cells were allowed to dry completely before staining them with 0.05% crystal violet for 10 minutes. Excess stain was removed by washing thoroughly with distilled water. Representative images were taken of the plates, after which cells were de-stained using 10% acetic acid for 15 minutes on a plate shaker, and the absorbance was measured at 590nm. For knockdown experiments, transfection of Silencer® Select siRNAs against CDK2 (Catalog # 4427038, ID: s204 and s206) was achieved using RNAiMax (ThermoFisher Scientific). For the validation of knockdown, cell lysate was collected after 72 hours of transfection for western blot analysis as described above.

### RT-PCR and expression profiling

RNA was isolated post-treatment using illustra RNAspin Mini RNA Isolation Kit (Cytiva) following the manufacturer instruction. The cDNA used for qPCR was prepared using qScript cDNA Synthesis Kit (Quanta Biosciences). Amplification was carried out as: 10 minutes at 95°C, 40 cycles of 30 seconds at 60°C, 30 seconds of extension, and a final extension of 5 minutes at 72°C. Primers are provided as a supplementary figure 1.

### Immunofluorescence

Cells were plated on glass slides and treated as indicated. For collection, media was removed and cells fixed with ice-cold 100% methanol and stored in -20°C. For staining, cells were washed with PBS three times, once with 5% BSA in PBS, blocked for 1 hour with 5% BSA in PBS, primary antibodies added (Anti-DNA-RNA Hybrid Antibody, clone S9.6, MERCK, MABE1095, p53BP1, Thermo Fisher Scientific, 83809-1-RR) for one hour incubation in room temperature, washing for 5minutes each with with 5% BSA in PBS was repeated thrice, secondary antibodies were added (Alexa Fluor™ 488 goat anti-mouse IgG, Alexa Fluor™ 546 goat anti-rabbit IgG) for one hour incubation in room temperature, washing for 5 minutes each with PBS and finally class slides were mounted with fluorescent mounting media (ProLong^™^ Glass Antifade Mountant with NucBlue^™^ Stain). Zeiss Axio Imager.Z2 upright epifluorescence wide-field microscope was used for imaging.

### Development of resistant cell line

To generate CDK2 inhibition-resistant cells, parental C4-2 cell line was cultured under BLU-222 using a stepwise dose-escalation strategy. Cells were initially exposed to 500 nM BLU-222 for two months followed by two months in 1000 nM, and finally, for additional two months in 1500 nM compound. Media was replaced every 48-72 hours.

### Statistical analysis and plots

All experiments were performed at least in triplicates and data are presented with standard error of mean. Statistical analyses and data visualization were performed using R studio. Statistical significance was determined using a paired sample two-tailed Student’s t-test, one-sample t-test and unpaired Mann-Whitney-Wilcoxon test, as appropriate and detailed when the data is presented, with p < 0.05 considered significant.

## Results

### Emergence of castration-resistant prostate cancer results in increased dependency on CDK2

We wanted to understand if development of CRPC is associated with altered expression of the key cyclins that control cell cycle progression. As detailed in the introduction, successful execution of cell cycle depends on the sustained phosphorylation of RB1, which releases E2F transcription factor to promote cell cycle. CDK4/6 together with the D-type cyclins (CCND) control decision to enter the cell cycle, while high activity of CDK2 brought about through the increased expression of CCNE is important to maintain the E2F activity throughout the cell cycle (**Fig. 1A**). After DNA synthesis is completed, the expression of CCNB1 increases, which stimulates CDK1 and completion of mitosis. We evaluated the expression of these major cell cycle cyclins when the normal prostate tissue transforms initially to primary prostate cancer, followed by establishment of metastasis and eventual castration-resistance. Strikingly, the expression of both the E-type cyclins (CCNE1 and CCNE2) was significantly increased while D-type cyclins (CCND1, CCND2 and CCND3) was significantly downregulated (**Fig. 1B**). The expression of the key regulator of mitosis, CDK1, and mitotic cyclins (CCNB1, CCNB2, CCNA1, CCNA2 and CCNA3) were also increased between normal prostate tissue and prostate cancer (**Suppl. Figs. 2A and 2B**). In contrast, the expressions of CDK2, CDK4 and CDK6 did not reveal robust changes between the tissue types (**Suppl. Fig. 2B**). Increased expression of CDK2, CCNE1 and CCNE2 was significantly associated with high Gleason Score (**Fig. 1C**). These data propose that prostate cancer cells, and particularly CRPC cells, are addicted to high activity of CDK2 to sustain the pro-proliferative phenotype.

**Figure 1.**
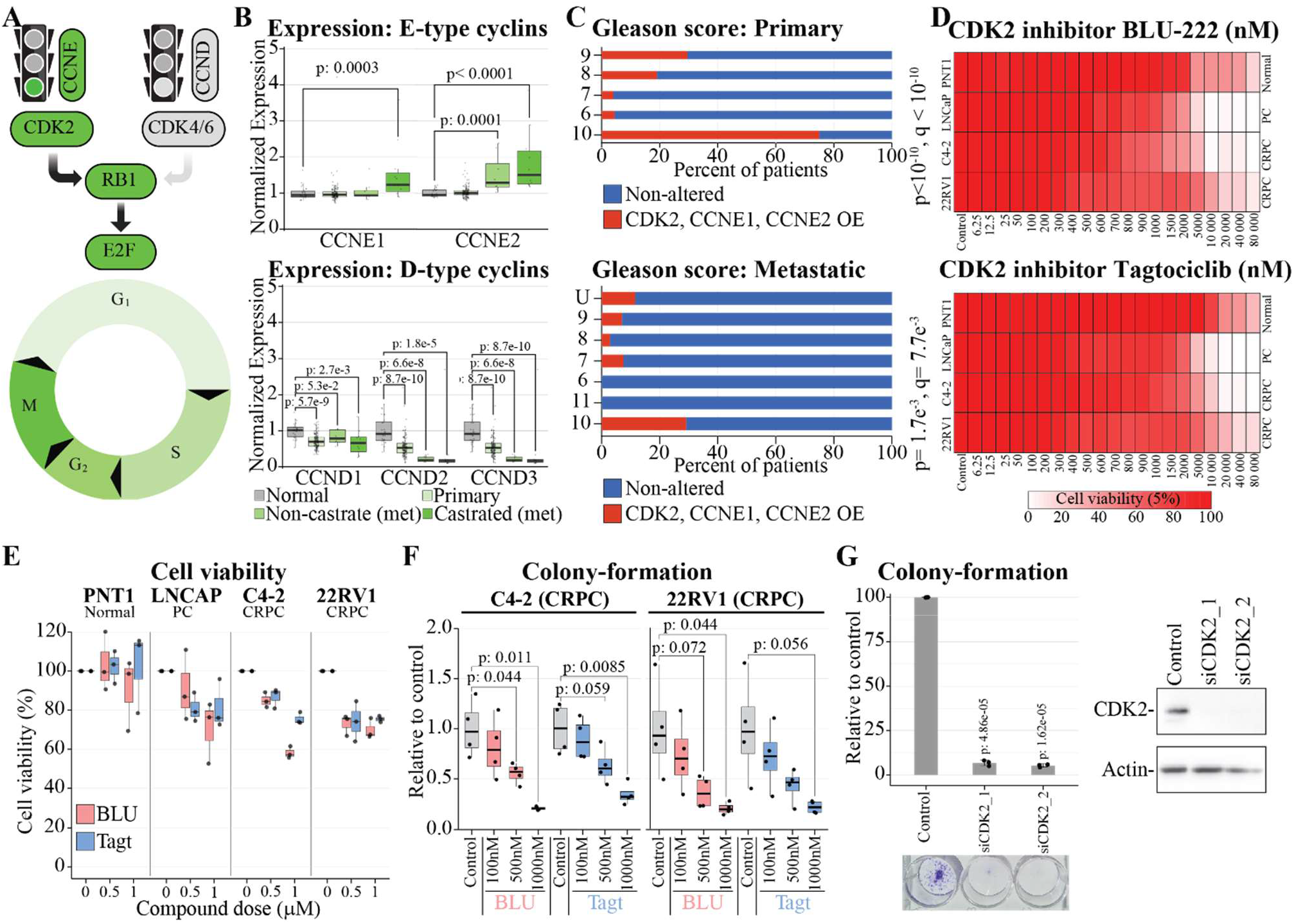
Development of aggressive, castration-resistant prostate cancer results in acquired dependency on CDK2. **A)** Schematic representation of RB1-E2F regulation during cell cycle. Phosphorylation of RB1 releases the E2F transcription factor, which activates the genes required for cell cycle progression. **B)** mRNA expression levels of E-type cyclins (*CCNE1* and *CCNE2*) and Dtype cyclins (*CCND1, CCND2*, and *CCND3*) were analyzed across normal prostate tissue, primary prostate cancer, metastatic prostate cancer, and castration-resistant prostate cancer using patient datasets obtained from [53] accessed through betastasis.com. Error bars represent the standard error of the mean (SEM) and unpaired Mann-Whitney-Wilcoxon test was used to evaluate statistical significance. **C and D)** Gleason score distribution in prostate tumors over-expression *CDK2* and / or *CCNE1* and / or *CCNE2* (exp > 2) accessed through the cBioPortal in the Prostate Adenocarcinoma (TCGA, Firehose Legacy) dataset and Metastatic Prostate Adenocarcinoma (SU2C/PCF Dream Team, PNAS 2019). Please, note that value of 11 is that recorded from cBioportal. **D and E)** Cell viability after 4 days of treatment with BLU-222 and Tagtociclib at the indicated concentrations (average of three biological replicates each with three technical replicates). Control samples were set to 100 and treatments were normalized to that. Error bars represent the SEM. **F and G)** Colony-formation assay after 7 days treatment with BLU-222 and Tagtociclib or CDK2 knockdown in C4-2 (3-4 biological replicates, paired two-tailed Student’s *t*-test **(F)** and one-sample *t*-test **(G)** were used to evaluate statistical significance and error bars represent the SEM. Western blot was used to confirm successful depletion of CDK2 after the knockdown (representative of three biological replicates).

We moved on to probe if depletion of CDK2 activity is toxic to prostate cancer, CRPC and a cell line derived from the normal prostate epithelia. First, we identified two selective and structurally divergent CDK2 inhibitors, BLU-222 and Tagtociclib [23, 24], and evaluated their efficacy against prostate cancer, CRPC and normal cells. As expected, both compounds suppressed proliferation of prostate cancer and CRPC cell lines with lower doses than they affected normal prostate cells (**Fig. 1D**). We identified 500 nM dose as a dose that has practically no effect on normal cells but suppressed proliferation of cancer cells by at least 20% (**Figs. 1D and 1E**). This dose also decreased the colony-formation ability of two CRPC models by 40-70% using either BLU-222 or Tagtociclib (**Fig. 1F and Suppl. Fig. 3**). Finally, we confirmed that knockdown of CDK2 leads to an almost complete loss of the colony formation ability of CRPC cells (**Fig. 1G**)

We have so far shown that transformation of normal prostate tissue to prostate cancer, and further to CRPC, is associated with transcriptional signature of CDK2-dependency and confirmed the acquired dependency using two distinct compounds. However, whilst we detected robust anti-proliferative effects, CRPC cells tolerated CDK2 inhibition reasonably well, and better understanding this should enable design of rational combinatorial therapies.

### Prolonged CDK2 inhibition transcriptionally upregulates CDK6-CCND2 axis

We developed CDK2 inhibitor-resistant cell line from the CRPC model C4-2 to establish if acquisition of resistance is explained by transcriptional rewiring of the CDK / cyclin-network. As expected, exposure of cells to CDK2 inhibitor suppressed proliferation and we therefore cultured the cells for an extended period of time in the presence of 500 nM BLU-222, a dose that had no effect on the normal prostate cells but suppressed proliferation of PC and CRPC cells (**Figs. 1D**). After 2 months, when cells had started to proliferate again, we increased the dose to 1000 nM for two months, followed up by 2 months in the presence of 1500 nM BLU-222. After this dose-escalation, our novel cell line tolerated a significantly higher dose of the BLU-222 compound (2-fold, **Fig. 2A**). They also became cross-resistant against the selective CDK2 inhibitor Tagtociclib but not against a pan-CDK inhibitor AT7519 [25] (**Figs. 2B and 2C**).

**Figure 2.**
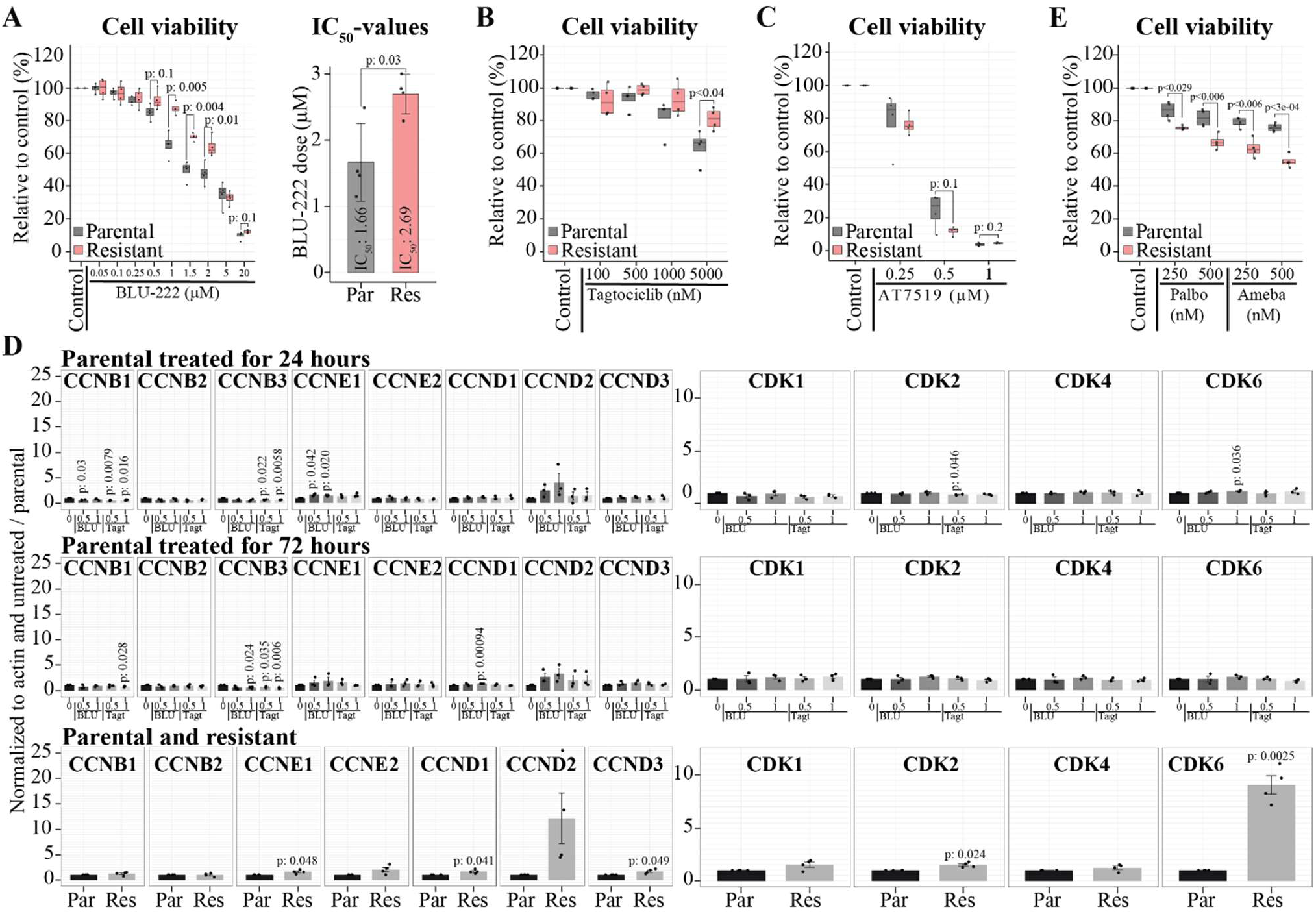
Resistance to CDK2 inhibition is associated with transcriptional rewiring to reinforce signalling through CDK6. **A)** Parental and BLU-222-resistant C4-2 cells were treated with BLU-222 as indicated and cell viability was assessed after 4 days of treatment (four biological replicates, each 6 technical replicates and paired two-tailed Student’s *t*-test was used to assess significance. GraphPad Prism was used to calculate IC_50_-values. Error bars represent the SEM. **B and C)** Parental and BLU-222-resistant C4-2 cells were treated with the selective CDK2 inhibitor Tagtociclib or the pan-CDK inhibitor AT7519 and viability assessed (the rest is the same as in 2A). **D)** RT-qPCR analysis of parental and BLU-222 resistant C4-2 cells after the indicated treatments. Data is from 3-4 biological replicates, error bars represent the SEM and statistical significance is evaluated by one sample *t*-test. **E)** Parental and BLU-222-resistant C4-2 cells were treated with the CDK4/6 inhibitor Abemaciclib and Palbociclib for 4 days and the rest is the same as in 2A.

We hypothesized that development of resistance occurs through transcriptional adaptation of the CDK / cyclin-network to compensate for the lower CDK2 activity. To assess this, we exposed the parental C4-2 cells to 500 nM BLU-222 and 500 nM Tagtociclib for 24 and 72 hours, isolated mRNA from untreated, treated and resistant cells, and evaluated the levels of the key cell cycle-cyclins. This experiment revealed a robust up-regulation of CCND2 at 24 hours (1.5 to 3-fold), 72 hours (1.5 to 3-fold) and in the resistant cells (10-fold) and also modest increase in the CCNE1 and CCNE2 mRNAs (**Fig. 2D**). At the same time, the expression of the mitotic cyclin CCNB decreased at both 24- and 72-hours and modestly increased in the resistant cells. These data propose that CRPC cells adapt to decline in CDK2 activity by up-regulating the CDK4/6 signaling. To further assess this, we evaluated the expression of the cell cycle kinases after short term and chronic CDK2 inhibition with BLU-222 and Tagtociclib. Indeed, the expression of CDK6 was upregulated by 9-fold in the resistant cells but this effect was not detected after 24- or 72-hours treatments (**Fig. 2D**). In addition, we noted a modestly increased expression of all of the other cell cycle kinases that positively regulate progression through different stages of the cell cycle in the cells exposed to CDK2 inhibitor chronically but not after 24- or 72-hours treatments (**Fig. 2D**).

Our transcriptional profiling data propose that acquisition of resistance to CDK2 inhibition causes acquired dependency on CDK6-signaling. To directly assess this, we treated our parental and resistant cells with CDK4/6 inhibitors Palbociclib [26] and Abemaciclib [27]. Indeed, the resistant cells were significantly more sensitive to both compounds (**Fig. 2E**).

We have so far shown that CRPC cells have acquired dependency on CDK2, which can be overcome through transcriptional adaptation of the CDK- / cyclin-network. This adaptation involves robust upregulation of the Cyclin D2 and CDK6, while the cell cycle progression is initially halted as evidenced by the decrease in mitotic cyclins. These conditions should lead to replication stress, which we moved on to probe next.

### CDK2 inhibition leads to formation of R loops and double-strand DNA breaks

CDK2 inhibition induced transcriptional changes to sustain cell cycle progression, and we hypothesized that these changes cause DNA replication stress. When transcription and replication machineries collide, this can lead to generation of R-loops, nucleic acid structures consisting of an RNA-DNA duplex and an unpaired DNA strand [28]. We assessed R-loop formation as the first port-of-call to evaluate CDK2 inhibition-induced replication stress. Indeed, treatment of C4-2 cells with either of the CDK2 inhibitors, BLU-222 or Tagtociclib, increased the R-loop levels as determined using immunofluorescence (**Fig. 3A**). We hypothesized that the accumulation of R-loops would lead to DNA damage. p53-binding protein 1 (p53BP1) is recruited to double strand breaks [29], and we used it as a marker for DNA damage here. Indeed, we noted a robust increase in the p53BP1-foci after treatment with both BLU-222 or Tagtociclib (**Fig. 3A**). Further inspection of the immunofluorescence data indicated that the R loops accumulate particularly in the densely packed areas in the nucleus, the areas responsible for synthesis of ribosomal RNA.

**Figure 3.**
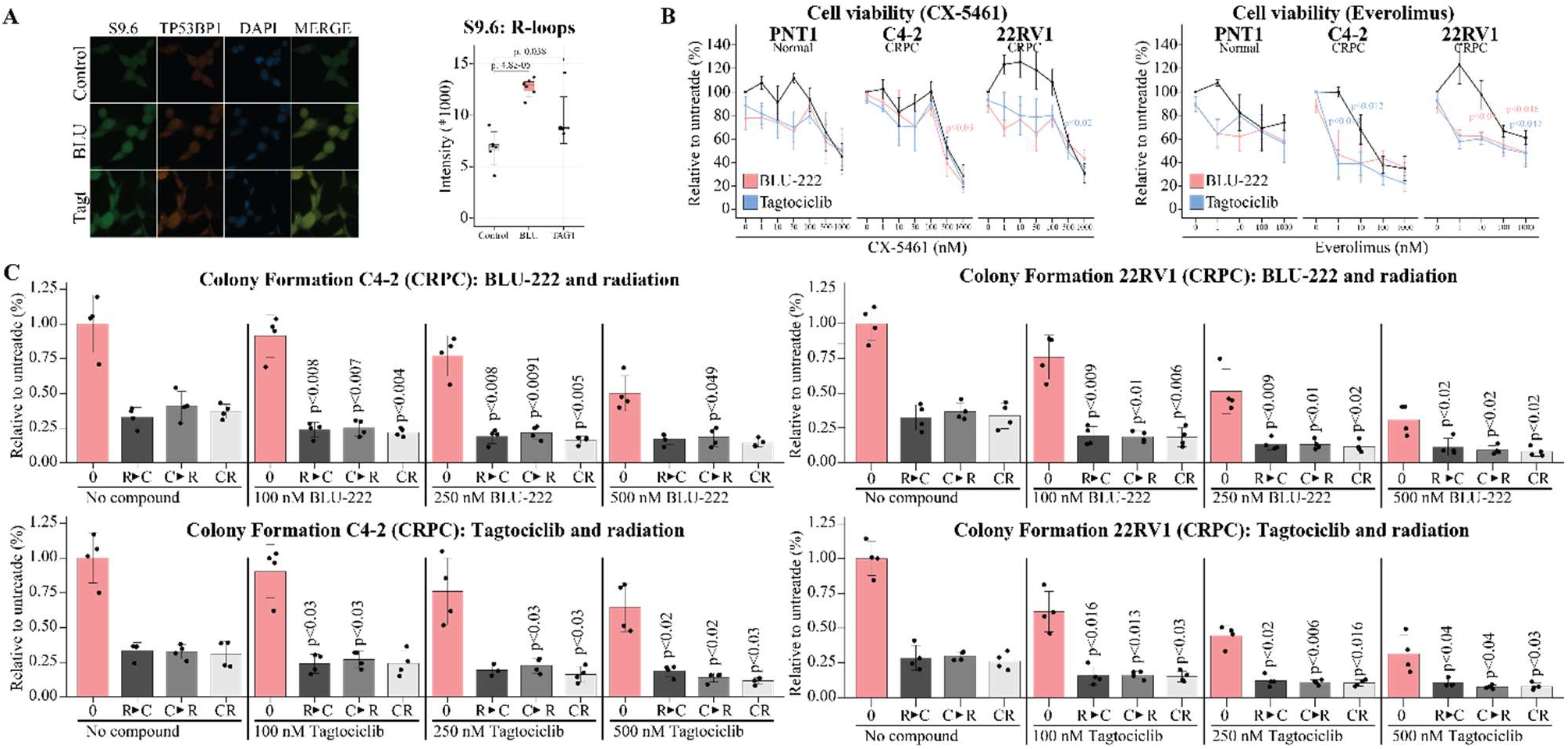
CDK2 inhibition causes replication stress and sensitizes prostate cancer cells to radiation. **A)** C4-2 cells were treated with BLU-222 (500 nM) and Tagtociclib (500 nM) for 3 days, R-loop accumulation was detected using immunofluorescence (the S9.6 antibody), and TP53BP1 was also detected. Quantification of signal intensity with SEM and significance was assessed by paired two-tailed Student’s *t*-test. **B)** CRPC cell lines (C4-2 and 22RV1) and normal prostate epithelial cells (PNT1) were treated for 4 days with CX-5461 or Everolimus alone, or in combination with BLU-222 (500 nM) or Tagtociclib (500 nM), cell viability assessed and data is the mean of four biological replicates (2-3 technical replicates each) with SEM. Paired two-tailed Student’s *t*-test was used to evaluate statistical significance, which is indicated in comparison between CDK2 inhibitor-alone and compound of interest-alone to combination and is color-coded according to CDK2 inhibitor used. **C)** Colony-formation assay. Cells were either not radiated (0), first radiated and then treated after 1 day (R>C), first treated and then radiated after 1 day (C>R) or treated and radiated at the same time (CR). After 7 days, the colonies were detected (4 biological replicates with SEM). Paired two-tailed Student’s *t*-test was used to evaluate statistical significance and the value reported is in comparison to both radiation-alone and CDK2 inhibitor-alone (whichever is higher, is reported).

We hypothesized that nucleolar stress and / or protein synthesis become points of vulnerability in combination with CDK2 inhibitors. For these experiments, we made use of CX-5461, a selective inhibitor of RNA Pol I, the polymerase responsible for ribosomal RNA synthesis in the nucleolus [30] and Everolimus, an inhibitor of the mTOR kinase, major regulator of protein synthesis [31]. According to our hypothesis, both CX-5461 and mTOR sensitized CRPC cells to CDK2 inhibitors, and the combination treatments were less toxic to a cell line derived from the normal prostate epithelia (**Fig. 3B**). However, the effect was modest, and at the best, additive, and different combination strategy is therefore required to effectively target the CRPC cells, and we moved on to acquire further evidence on the DNA damage-activation in response to CDK2 inhibition.

In support of activation of the DNA damage response, we detected robust accumulation of p53 as determined by western blotting after CDK2 knockdown, or treatment with BLU-222 and Tagtociclib (**Suppl. Fig. 4**). p53 functions as an important mediator of the cellular response to DNA damage [32]. However, we earlier detected a clear decrease in the proliferation of CRPC cells that either have wild type p53 (C4-2) or mutated one (22RV1, **Figs. 1D, 1E and 1F**). Therefore, p53 itself is not necessary for the CDK2 inhibition-induced anti-proliferative effects.

We hypothesized that CDK2 inhibition-induced anti-proliferative effects could be significantly increased if DNA damage was augmented. In the clinical setting, prostate cancer can be treated by radiotherapy, which effectively suppresses the tumor progression. However, radiation should arrest the cell cycle, which might decrease the efficacy of CDK2 inhibitors. To take this into account, we devised a four-armed experimental strategy: single agent, first radiotherapy followed by CDK2 inhibitor, first CDK2 inhibitor followed by radiotherapy, and simultaneous treatment. Strikingly, all of these combinatorial treatment strategies significantly further suppressed the colony-formation ability of both C4-2 and 22RV1 when compared to single agent treatments (**Fig. 3C and Suppl. Fig. 5**).

We have so far shown that CDK2 inhibition causes modest anti-proliferative effects but prominent DNA replication stress, which significantly sensitizes CRPC cells to radiotherapy. Next, we moved on to systematically probe if specific clinically utilized anti-CRPC therapies could be combined with CDK2 inhibitors to cause cancer cell-selective anti-proliferative effects.

### CDK2 inhibition results in increased dependency on the high activity of androgen receptor

We established a library of compounds from currently used prostate cancer therapies to establish which would form the most effective combination with CDK2 inhibitors. The compounds in our library can be divided in two major groups: First, we have five cytotoxic compounds and second, more targeted compounds. In the first class we included Etoposide, which targets topoisomerase II enzyme and thereby leads to defective resolution of negative and positive supercoils in DNA [33]. Carboplatin and Cisplatin cause inter- and intra-DNA adducts, which interfere with DNA replication and transcription [34]. Docetaxel and Cabazitaxel target the assembly of microtubules into the mitotic spindle thereby triggering G2/M arrest [35]. For more biomarker-guided and disease-specific therapies, we included Olaparib, Rucaparib, Enzalutamide and Darolutamide. Olaparib and Rucaparib target PARP, which becomes a synthetic point of lethality in cells that have mutations in homologous recombination pathway [36], while Enzalutamide and Darolutamide are inhibitors of androgen receptor [37]. To identify combination(s) that are less toxic to normal cells, we included a cell line derived from normal prostate epithelia, PNT1, and two models of CRPC, C4-2 and 22RV1. The screen was performed across four biological replicates and 2-3 technical replicates in each plate to assure robustness.

Interestingly, the efficacy of CDK2 inhibitors was greatly augmented by cytotoxic-compounds, while the targeted therapies had more variable efficacy (**Fig. 4A**). In more detail, Docetaxel, Cabazitaxel, Cisplatin, and Carboplatin were toxic as single agents to all the three model systems and CDK2 inhibition predominantly increased this toxicity. In contrast, PARP inhibitors and Etoposide were toxic at the single agent level to all of the cell line models, but did not clearly enhance the efficacy of CDK2 inhibitors (**Fig. 4B**). Interestingly, both anti-androgens, Enzalutamide and Darolutamide, significantly sensitized both of the CRPC cell lines to both of the CDK2 inhibitor (**Fig. 4C**). Importantly, anti-androgens were not toxic to the normal PNT1 cells and did not alter the efficacy of CDK2 inhibition in the normal cells (**Fig. 4C**). To acquire further confidence on the fact that prostate cancer cells, but not normal cells, are selectively sensitive to combined targeting of CDK2 and androgen receptor, we repeated the last treatment combination in LNCaP cells, which represent androgen dependent prostate cancer and RWPE-1, a cell line representing normal prostate epithelia. Indeed, we detected a highly significant combinatorial toxicity selectively in LNCaP cells but not in RWPE-1 cells (**Fig. 4D)**. Colony-formation assays revealed that combination of either anti-androgens or complete hormone-depletion from the media significantly sensitize both C4-2 and 22RV1 CRPC cells to CDK2 inhibitor BLU-222 (**Fig. 4E and Suppl. Fig. 6**). Finally, we confirmed the importance of high CDK2 levels and activity for CRPC cell response against anti-androgens using knockdown experiments and colony-formation assays with both anti-androgens and complete removal of hormones from the media (**Suppl. Fig. 7**). These data identify the androgen receptor as a point of vulnerability in combination with CDK2 inhibitors.

**Figure 4.**
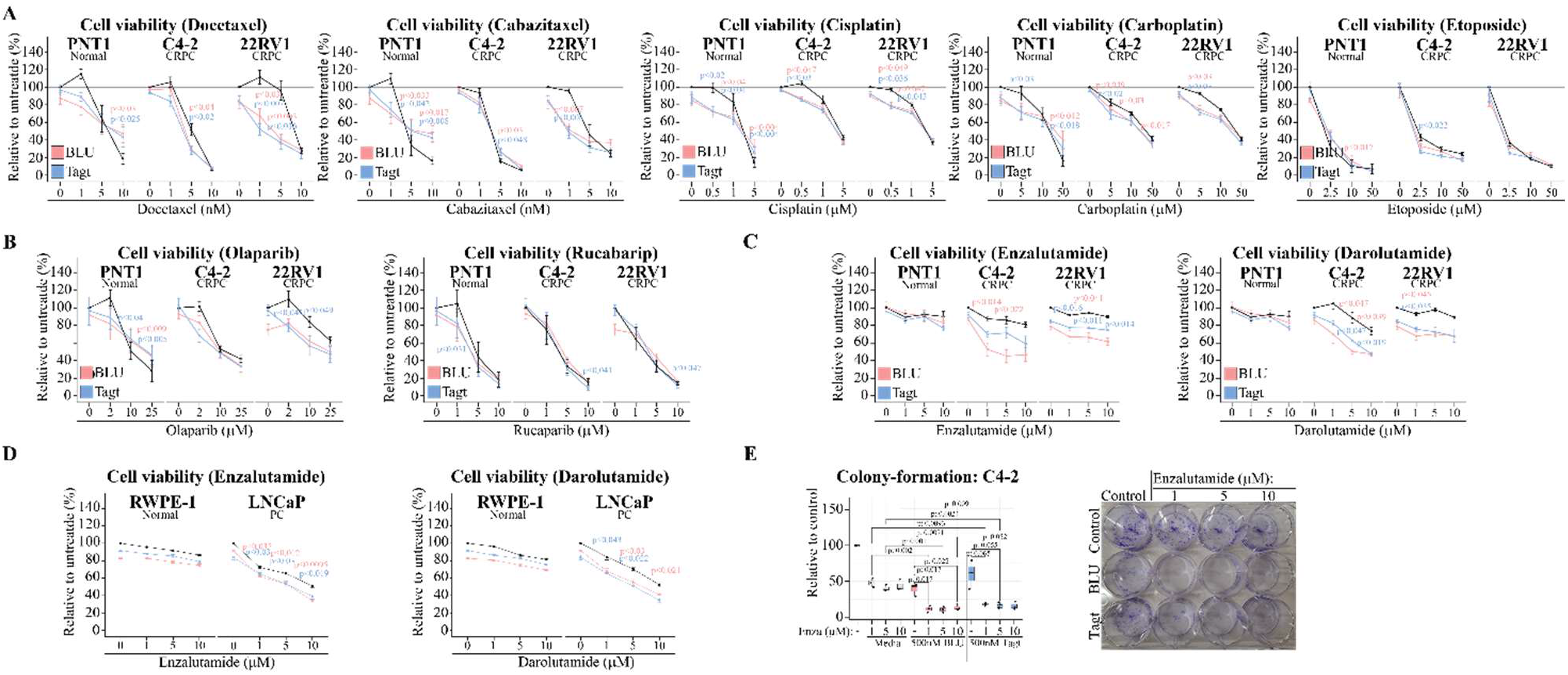
Targeted compound screen reveals that CDK2 inhibitors re-sensitize castration-resistant prostate cancer cells to anti-androgens. **A-D)** CRPC cell lines (C4-2 and 22Rv1), cell lines derived from normal prostate epithelia (PNT1 and RWPE-1), and prostate cancer cells (LNCaP) were treated for 4 days as indicated (BLU-222: 500 nM and Tagtociclib: 500 nM), viability assay performed and data is the mean of four biological replicates (2-3 technical replicates each) with SEM. Paired two-tailed Student’s *t*-test was used to evaluate statistical significance, which is indicated in comparison between CDK2 inhibitor-alone and compound of interest-alone to combination and is color-coded according to CDK2 inhibitor used. **E)** Colony-formation assay. Cells were treated as indicated and colony-formation assay performed after 7 days (3 biological replicates with SEM and paired two-tailed Student’s *t*-test was used to assess the significance).

### CDK2 inhibition stimulates androgen receptor activity

Next, we wanted to understand if CDK2 inhibition alters AR levels and / or activity acutely and / or chronically. Interestingly, CDK2 inhibition stimulated AR activity already after short-term treatment (24 hours) in both of the CRPC models assessed (C4-2 and 22RV1) as determined by the expression of AR target genes *KLK3* and *GAPDH*, which were increased up to 4-fold, but the levels of AR remained unchanged (**Fig. 5A**). We observed a similar response in our CDK2 inhibitor-resistant cell line revealing a 3-fold increase in KLK3 (**Fig. 5B**). We wanted to understand if altered CDK2 activity is associated with altered AR-signaling also in patient tumors and used the metastatic prostate cancer sample set available through the cBioPortal to assess this. Interestingly, increased expression of CDK2 and / or its cyclins was significantly associated with the neuroendrocrine-phenotype (**Fig. 5C**). These data show that CDK2 inhibition increases androgen receptor activity in CRPC cells, which can be therapeutically targeted using anti-androgens.

**Figure 5.**
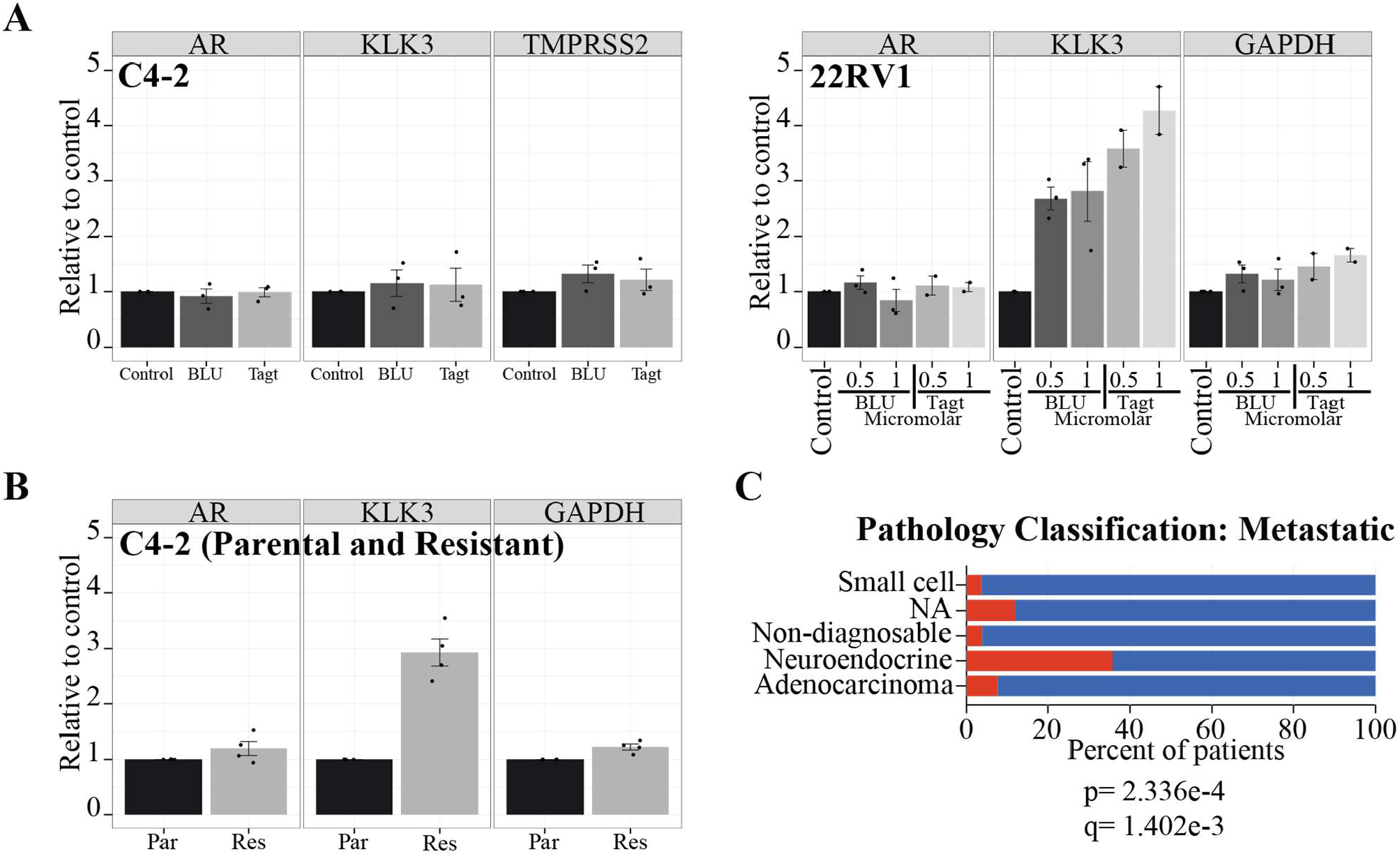
Adaptation to CDK2 inhibition stimulates androgen receptor activity acutely and chronically. **A)** CRPC cell lines C4-2 and 22Rv1 were treated with BLU-222 or Tagtociclib for 48 hours. mRNA levels of AR and AR target genes were measured by qPCR. For C4-2 cells, AR, KLK3, and TMPRSS2 expression were analyzed, whereas for 22RV1 cells, AR, KLK3, and GAPDH expression were analyzed. Boxplots represent data from 2-3 biological replicates. Error bars represent the SEM. **B)** mRNA levels of AR and AR target genes, KLK3, and GAPDH were quantified by qPCR in parental and BLU-222-resistant C4-2 cells. Data is from four biological replicates with SEM. **C)** Pathology classification of prostate tumors over-expressing *CDK2* and / or *CCNE1* and / or *CCNE2* (exp > 2) accessed through the cBioPortal in the Metastatic Prostate Adenocarcinoma (SU2C/PCF Dream Team, PNAS 2019).

In brief, here we show that development of prostate cancer is associated with increased dependency on CDK2, which becomes an actionable point of vulnerability as cells develop CRPC phenotype. We anticipate that appropriate sequencing of either targeted therapies or cytotoxic compounds is required for efficacy in the future clinical trials with CDK2 inhibitors against prostate cancer.

## Discussion

Dysregulation of cell cycle enables cancer cells to proliferate rapidly but may also open therapeutic vulnerabilities, which has prompted development of targeted therapies. Here, we show that development of CRPC leads to acquired dependency on CDK2 while the role of CDKs4/6 diminish in parallel. Using two selective CDK2 inhibitors in clinical development along with knockdown strategies we reveal that CDK2 forms a combinatorial lethal target with the existing prostate cancer therapeutics, most notably radiotherapy and anti-androgens.

The cyclic regulation of specific cyclins drives the initiation, execution, and termination of the cell cycle, and cancer cells appear to augment signalling through a specific cyclin-CDK pairs. Here, we show that development of CRPC is associated with an increased expression of mitotic cyclins, which likely represents a higher proportion of actively cycling cells within the tumour rather than overall increased expression in only a few cells (**Suppl. Fig. 2**). In fact, there are clinically utilized tests that measure the expression of cell cycle-mRNAs and predict how aggressive the patient’s tumor is, such as the Prolaris Score [38]. Strikingly, however, we show here that the CRPC cells downregulate CCNDs and upregulate CCNEs, implying a selective dependency on CDK2 (**Fig. 1B**).

We show that CRPC cells are dependent on CDK2, which we propose is explained by the underlying factor driving the cell cycle progression. We show that both hormone-dependent and CRPC cells are sensitive to CDK2 inhibitors BLU-222 and Tagtociclib (**Figs. 1D, 1E and 1F**). Earlier, despite the initial positive results using CDK4/6 inhibitors as a prostate cancer therapy, these compounds have been discontinued due to lack of efficacy [17, 39]. This is likely explained by the fact that in prostate cancer multiple growth stimuli synergize to promote cell cycle, in particular androgens and epidermal growth factor, which activate CDK2 but not CDK4 or CDK6 [40]. Androgen receptor drives the expression of a large number of cell cycle regulators but, curiously, does not prominently affect the expression CDK4/6 or their cyclins [41]. Instead, it acts by downregulating the endogenous inhibitor of both CDK2 and CDK4, p27 [40, 42]. Accordingly, over-expression of G1/S cyclins is not sufficient to confer androgen-independent proliferation [43]. In a striking contrast to this, the hormone positive breast cancer cells drive the expression of *CCND*-gene through a tissue-specific enhancer [44]. Accordingly, these cells are particularly sensitive to compounds targeting CDK4/6, as initially demonstrated through *in vitro* and animal experiments [13, 26], and later on in clinical trials and eventual approval as therapy [45, 46]. However, emergence of resistance remains an issue in the case of CDK4/6 inhibitors [45], as is likely the case for CDK2 inhibitors based on the data presented in this manuscript (**Fig. 2**). Importantly, however, we identify rational strategies to circumvent the emerging resistance and augment the efficacy of CDK2 inhibitor-based therapy.

We show that CDK2 inhibition triggers transcription-replication conflicts in prostate cancer cells, which opens a combinatorial treatment strategy. Using immunofluorescence, we demonstrated accumulation of R-loops, particularly in the densely-packed regions of the nucleus, likely representing nucleoli where ribosomal RNA is synthesized (**Figs. 3A and 3B**). This prompted us to ask if CDK2 inhibitors sensitize cells to compounds interfering directly with ribosomal RNA synthesis through RNA Pol I inhibitors or translation by targeting mTOR; however, whilst the effects were significant, they were rather modest (**Fig. 3C**). Nevertheless, we noted robust accumulation of R-loops, which are known to activate the innate immune response [47-49]. In the future, it is therefore of a high interest to establish if CDK2 inhibitors can augment anti-tumor immunity, as this could lead to curative therapy. In support of this, we show that radiotherapy greatly augments the anti-proliferative effect of CDK2 inhibition (**Fig. 3D**).

We developed a CDK2-inhibitor resistant cell line and performed targeted combinatorial lethality screen to understand how these cells escape from the initially cell cycle-halting therapy (**Figs. 3, 4A, 4B and 4C**). The experience from treating breast cancer patients with CDK4/6 inhibitors have revealed escape mechanisms particularly through CDK2 and RB1, which provide rational targeted therapy with the recently developed CDK2 inhibitors [50, 51]. In support of this, we show that resistance to CDK2 inhibition sensitizes these cells to compounds targeting CDK4/6 (**Fig. 2E**). However, current clinical trials assessing the CDK2 and CDK4/6 inhibition combinations are either terminated with no data disclosed (NCT05252416) or are not actively recruiting (NCT04553133). In addition, combination of CDK2 and CDK4/6 inhibitors, whilst likely effective against cancer cells, can be expected to result in on-target toxicity due to effects on normal cells. Therefore, we set up a library of the therapies currently used against prostate cancer, which revealed cancer cell-selective sensitization to anti-androgens (**Figs. 4A, 4B and 4C**). Interestingly, CDK2 inhibition also robustly stimulated AR activity (**Figs. 5A and 5B**). These data suggest that inhibiting CDK2 restores sensitivity to anti-androgen therapy in CRPC cells that were previously resistant. Curiously, this is the same principal rational as utilized against breast cancer where the anti-estrogen receptor therapy is successfully combined with CDK4/6 inhibitors [52]. Overall, both the existing literature and the data presented in this manuscript underscore the importance of the underlying tissue-specific transcriptional program as a targetable vulnerability in combination with CDK-inhibition.

## Conclusions

In brief, we show that prostate cancer cells upregulate E-type cyclins (CCNE1/2) and are highly dependent on CDK2. Our data provide rational for combinatorial treatment strategy with existing therapies, in particular anti-androgens and radiotherapy together with CDK2 inhibitors.

## List of abbreviations

AR: androgen receptor
ADT: androgen deprivation therapy
CDK: cyclin dependent kinase
CCNA: cyclin A
CCNB: cyclin B
CCND: cyclin D
CCNE: cyclin E
CRPC: castration resistant prostate cancer
PC: prostate cancer
RB1: Retinoblastoma protein 1

## Declaration

### Ethics approval and consent to participate

Not applicable.

### Consent for publication

Not applicable.

### Availability of data and materials

All the data and materials are presented as part of this manuscript.

### Authors’ contributions

JC evaluated the expression of factors of interest in patient samples, performed the combinatorial screen with prostate cancer-relevant therapies, generated all figures in R and wrote the initial draft of introduction, discussion, and figure legends. AM performed significant part of the wet lab experiments. SY assisted AM in experiments and contributed to manuscript writing. HMI conceptualized the study, acquired resources and wrote the initial draft of the manuscript. All authors provided critical feedback on the manuscript.

## Acknowledgements

Imaging was performed at the Biomedicum Imaging Unit, the University of Helsinki, supported by the Helsinki Institute of Life Science (HiLIFE) and Biocenter Finland.

