## Supplementary figures and supplementary figure legends for "Castration-resistant prostate cancer cells are addicted to the high activity of cyclin-dependent kinase 2"

| Primer | Sequence |
|---|---|
| CCNE1_F | AAGGAGCGGGACACCATGA |
| CCNE1_R | ACGGTCACGTTTGCCTTCC |
| CCNE2_F | TCAAGACGAAGTAGCCGTTTAC |
| CCNE2_R | TGACATCCTGGGTAGTTTTCCTC |
| CCND1_F | GCTGCGAAGTGGAAACCATC |
| CCND1_R | CCTCCTTCTGCACACATTTGAA |
| CCND2_F | ACCTTCCGCAGTGCTCCTA |
| CCND2_R | CCCAGCCAAGAAACGGTCC |
| CCND3_F | TACCCGCCATCCATGATCG |
| CCND3_R | AGGCAGTCCACTTCAGTGC |
| CCNB1_F | AATAAGGCGAAGATCAACATGGC |
| CCNB1_R | TTTGTTACCAATGTCCCCAAGAG |
| CCNB2_F | CCGACGGTGTCCAGTGATTT |
| CCNB2_R | TGTTGTTTTGGTGGGTTGAACT |
| CCNB3_F | ATGAAGGCAGTATGCAAGAAGG |
| CCNB3_R | CATCCACACGAGGTGAGTTGT |
| CCNA1_F | GAGGTCCCGATGCTTGTCAG |
| CCNA1_R | GTTAGCAGCCCTAGCACTGTC |
| CCNA2_F | CGCTGGCGGTACTGAAGTC |
| CCNA2_R | GAGGAACGGTGACATGCTCAT |
| CDK1_F | AAACTACAGGTCAAGTGGTAGCC |
| CDK1_R | TCCTGCATAAGCACATCCTGA |
| CDK2_F | CCAGGAGTTACTTCTATGCCTGA |
| CDK2_R | TTCATCCAGGGGAGGTACAAC |
| CDK4_F | ATGGCTACCTCTCGATATGAGC |
| CDK4_R | CATTGGGGACTCTCACACTCT |
| CDK6_F | GCTGACCAGCAGTACGAATG |
| CDK6_R | GCACACATCAAACAACCTGACC |
| AR_F | TCGACCATTTCTGACAACGC |
| AR_R | GAAGCTGTTCCCCTGGACTC |
| KLK3_F | GCAGCATTGAACCAGAGGAG |
| KLK3_R | AGAACTGGGGAGGCTTGAGT |
| TMPRSS2_F | GTCCCCACTGTCTACGAGGT |
| TMPRSS2_R | CAGACGACGGGGTTGGAAG |
| GAPDH_F | ATCTTCTTTTGCGTCGCCAG |
| GAPDH_R | CGAACACATCCGGCCTGC |
| ACTIN_For | TGGGACGACATGGAGAAAAT |
| ACTIN_Rev | AGAGGCGTACAGGGATAGCA |

### Supplementary figure 1. Primers used in this study.

**A**

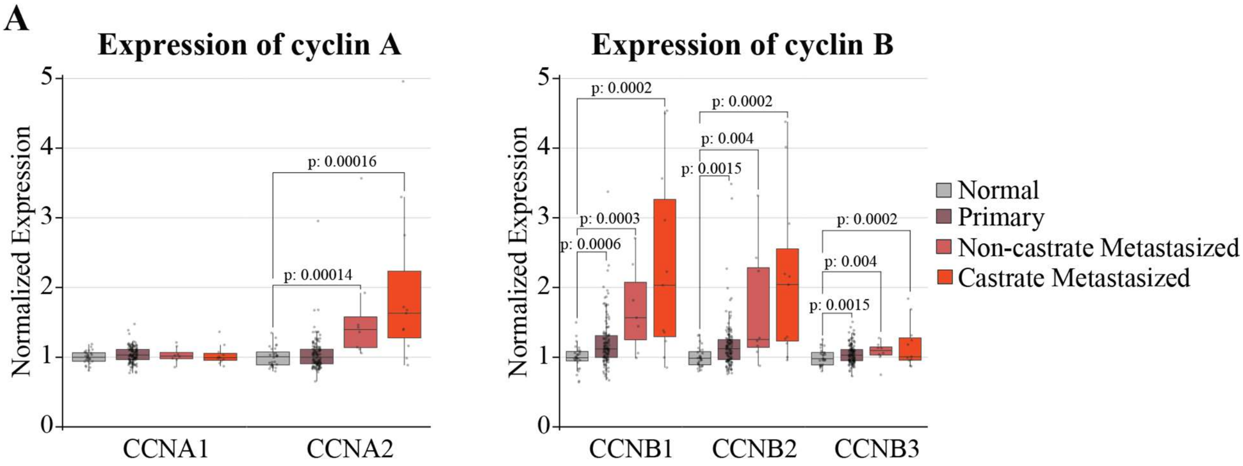


**B**

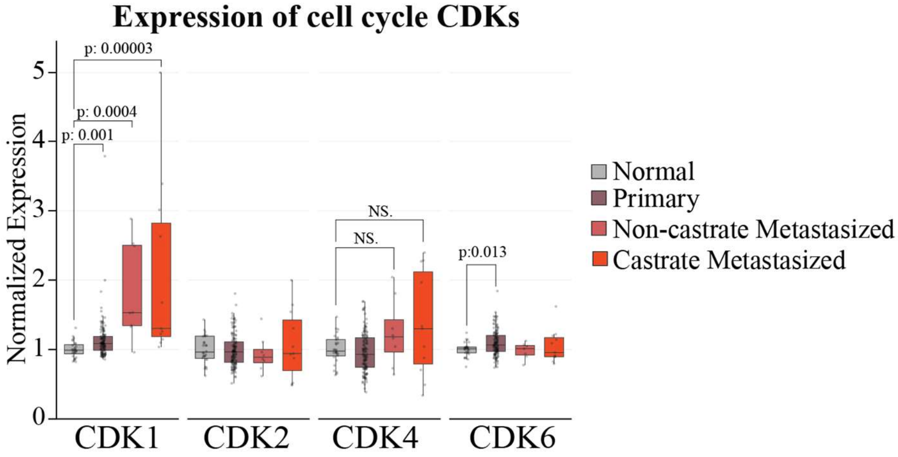


**Supplementary figure 2. Mitotic cyclins and CDK1 are over-expressed in primary and metastatic prostate cancer.** mRNA expression levels of cyclins and cyclin-dependent kinases were analyzed across normal prostate tissue, primary prostate cancer, metastatic prostate cancer, and castration-resistant prostate cancer using patient data obtained from [53] accessed through betastasis.com. Error bars represent the standard error of the mean (SEM) and unpaired Mann-Whitney-Wilcoxon test was used to evaluate statistical significance.

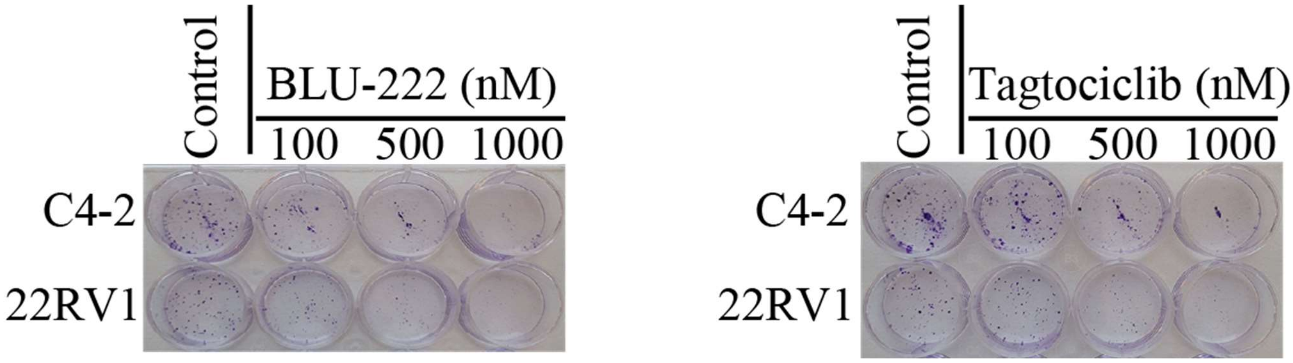


**Supplementary figure 3. Examples images of colony-formation experiment.** Colony-formation assay after 7 days treatment with BLU-222 and Tagtociclib or CDK2 knockdown (4 biological replicates).

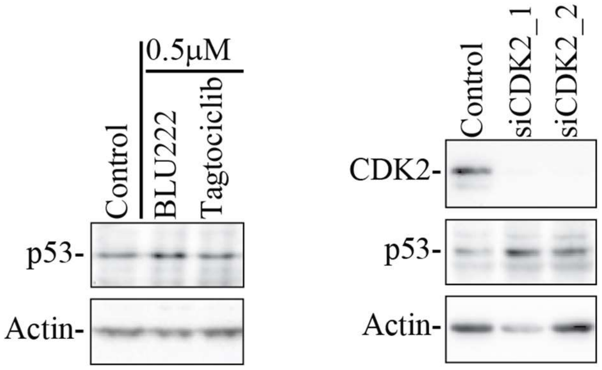


**Supplementary figure 4. CDK2 inhibition or knockdown increases p53 levels.** C4-2 cells were treated as indicated or transfected with CDK2-targeting siRNAs, and protein levels of CDK2, p53 and actin were analyzed by western blotting. Data is representative of three biological replicates.

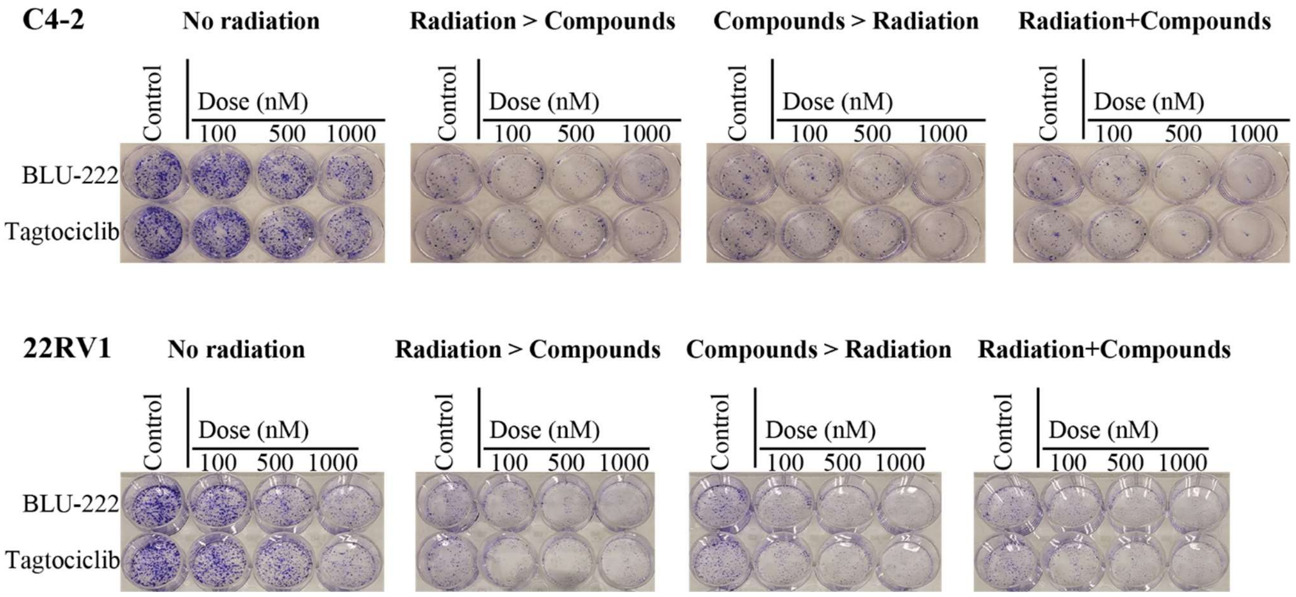


**Supplementary figure 5. Examples images of colony-formation experiment.** Colony-formation assay. Cells were either not radiated (C), first radiated and treated after 1 day (R>C), first treated and then radiated after 1 day or treated or radiated and treated at the same time (CR). After 7 days, the colonies were detected. This is representative of 4 biological replicates.

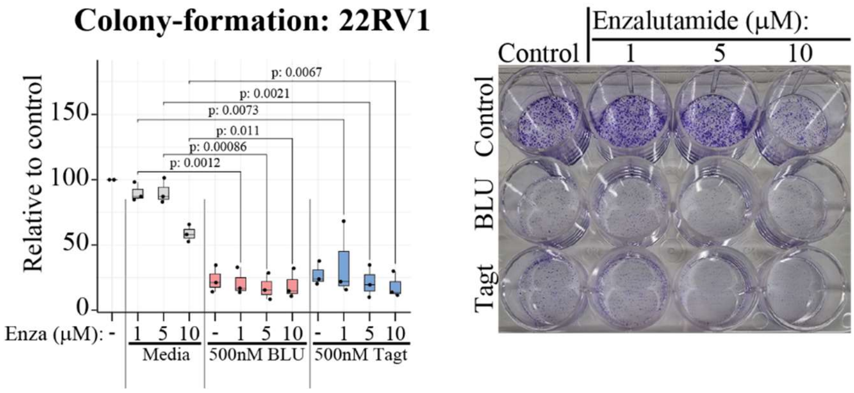


**Supplementary figure 6. CDK2 inhibition sensitizes CRPC cells to anti-androgen Enzalutamide.** 22RV1 cells were treated as indicated and colony-formation assay performed after 7 days (3 biological replicates with SEM and two-tailed Student's *t*-test was used to assess the significance).

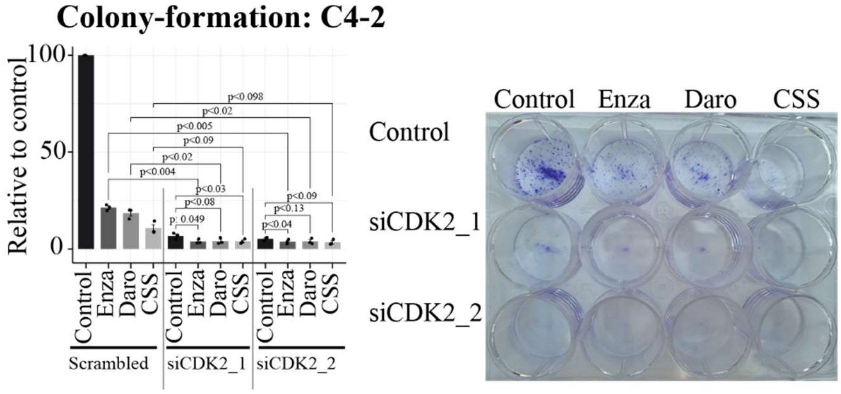


**Supplementary figure 7. Knockdown of CDK2 sensitizes castration-resistant prostate cancer cells to anti-androgens.** CDK2 was knocked down for a day after which C4-2 cells were treated as indicated and colony-formation assay performed after 7 days (3 biological replicates with SEM and paired samples two-tailed Student's t-test was used to assess the significance). Darolutamide and Enzalutamide dose: 5μM, CSS= androgen starvation (charcoal-stripped serum).
